# Region-Specific Transcriptional Signatures of Brain Aging in the Absence of Neuropathology at the Single-cell Level

**DOI:** 10.1101/2023.07.31.551097

**Authors:** Monica E. Mesecar, Megan F. Duffy, Dominic J. Acri, Jinhui Ding, Rebekah G. Langston, Syed I. Shah, Mike A. Nalls, Xylena Reed, Sonja W. Scholz, D. Thad Whitaker, Pavan K. Auluck, Stefano Marenco, Alex R. DeCasien, J. Raphael Gibbs, Mark R. Cookson

## Abstract

Given that age is a significant risk factor for multiple neurodegenerative diseases, investigating normal brain aging may help identify molecular events that may contribute to increased disease risk over time. Single-nucleus RNA sequencing (snRNA-seq) enables analysis of gene expression changes within specific cell-types, potentially offering insights into the molecular mechanisms underlying aging. However, most brain snRNA-Seq datasets used age-matched controls from studies focused on pathological processes and have largely been limited to cortical regions. Therefore, there is a need to investigate the non-pathological aging process in brain regions that are vulnerable to age-related diseases. Here, we report a snRNA-seq study of 6 young (20–30 years) and 7 aged (60–85 years) encompassing four different brain regions: the entorhinal cortex, middle temporal gyrus, subventricular zone, and putamen. We captured over 150,000 nuclei that represented 10 broad cell-types. While we did not find statistically significant differences in cell-type proportions with age, region- and cell-type-specific differential expression analyses identified over 8,000 age-associated genes. Notably, within a given cell-type, most of these associations were region-specific. Functional enrichment analyses of the gene sets for each cell-type-region combination revealed diverse biological processes, including multiple hallmarks of aging, such as proteostasis, interactions with cytokines, vesicular trafficking, metabolism, inflammation, and metal ion homeostasis. Overall, our findings suggest that unique cell-types exhibit distinct transcriptional aging profiles both at the cell-type level and across different brain regions.

## Introduction

Age is a common risk factor for multiple neurodegenerative diseases (NDDs), including Alzheimer’s disease, Parkinson’s disease, and amyotrophic lateral sclerosis (ALS)^1^. These age-related neurodegenerative diseases each have patterns of regional susceptibility. However, it remains unclear whether these regional differences are intrinsically linked to disease-causing mechanisms or reflective of global changes in brain health as an individual ages. Studies of NDDs often use age-matched controls without similar neuropathology to control for age as a source of variability. However, there is still a critical need to understand which cell-type- and region-specific effects arise specifically in individuals without NDDs during normal brain aging.

Prior studies have used bulk transcriptomic and epigenomic analyses to explore the effects of aging on the human brain. Major themes emerging from these investigations include loss of synaptic gene expression and acquisition of inflammatory signaling networks^2^. Our previous RNA-sequencing (RNA-seq) analyses of the human dorsolateral prefrontal cortex (dlPFC) identified networks of gene expression changes with age, including loss of neuronal genes^3^. For example, there was a strong decrease in *SST*, the gene encoding the peptide neurotransmitter somatostatin, which defines one of the major subtypes of inhibitory neurons in the human cerebral cortex.

Although the literature suggests that robust changes in gene expression correlate with age, several aspects of the data remain difficult to interpret. For example, loss of neuronal markers could reflect a loss of neurons or changes in gene expression among neurons. Recent development of single-cell methods allows for the identification of unique cell-types and the interrogation of gene expression within each population. However, to date, single-cell studies of postmortem human brain tissue have focused primarily on the cerebral cortex^4–6^. Therefore, it is unknown whether cell-type-specific aging signatures are similar across different brain regions. Given the known effects of non-pathological aging on the human cerebral cortex, we aimed to investigate which age-effects were regionally distinct versus shared broadly across regions that have shown vulnerability to age-related diseases.

We included two cortical areas that are susceptible to Alzheimer’s disease (AD) pathology: the entorhinal cortex (EC), which is affected early in the disease, and the middle temporal gyrus (MTG), which is affected at later stages^7,8^. We also included the putamen (PUT), which is affected by Huntington’s disease (HD)^9,10^ and is the target of dopaminergic neurons that are lost in Parkinson’s disease (PD)^11–13^ and the subventricular zone (SVZ) region, which is permissive for neurogenesis during development, although whether this remains true in the adult human brain is still contested^14–16^. By curating a cohort of young (20–30 years old) and aged (60–85 years old) individuals who lacked evidence for NDD-related pathology, we aimed to document age-related transcriptional changes across different cell-types and regions. We did not observe statistically significant differences in cell-type proportions across age with this sample size. However, in contrast, we did find large effects of age on gene expression of specific genes per cell-type. Notably, the vast majority of age-dependent differentially expressed genes (aDEGs) were unique to specific regions or cell-types. We find that aDEGs within specific cell-type–region combinations are associated with previously reported hallmarks of aging^17,18^, including proteostasis, interactions with cytokines, vesicular-trafficking, metabolism, the immune system and inflammation, and metal ion homeostasis. Taken together, our data serve as a resource for those studying age-related processes in the absence of disease-associated neuropathology.

## Results

### Cell-type-specific profiles of aging across multiple human brain regions

We aimed to investigate how gene expression profiles of different brain cell populations change with age across distinct brain regions. To this end, postmortem human brain tissue samples were obtained from 4 brain regions, including the entorhinal cortex (EC), middle temporal gyrus (MTG), subventricular zone (SVZ), and putamen (PUT). For 13 individuals, including six younger (20–30 years old) and seven older (60–85 years old) donors, 12 tissue samples were obtained for each of the four regions for a final sample of 48 samples. Donor groups were balanced for sex and tissue number (see Supp. Fig. 1 and Supp. Table 1 for donor and tissue demographics). Nuclei were isolated from the selected tissue samples, processed, and sequenced as described in Methods and shown schematically in Fig.1A.

**Figure 1:**
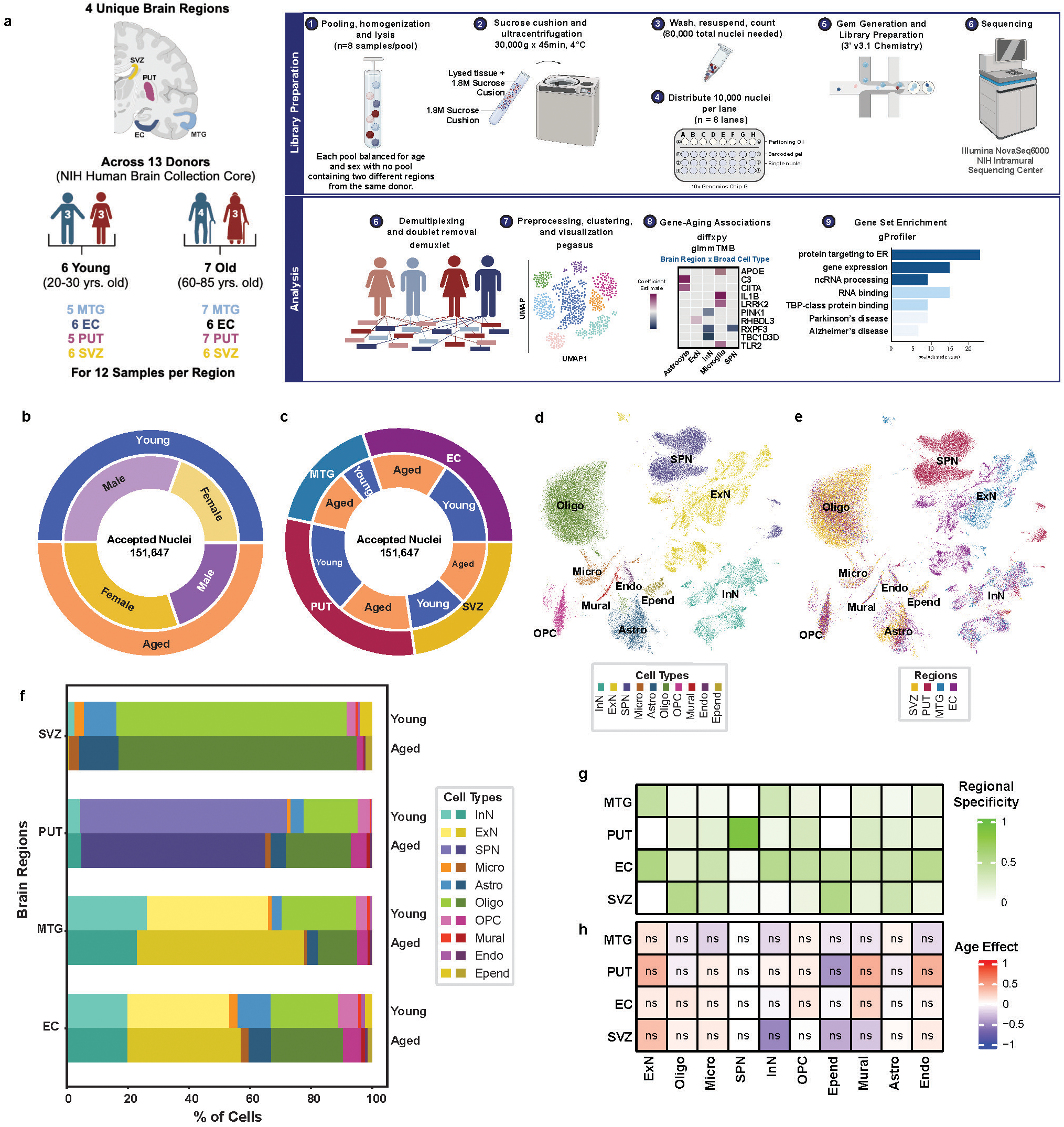
A single cell dataset for human brain aging across cell types and regions. **(a)** An outline of the single-nucleus RNA sequencing study design, including sample selection and workflow for investigating age-related differentially expressed genes (aDEGs) across 4 human brain regions––Entorhinal cortex (EC), Middle Temporal Gyrus (MTG), Putamen (PUT), and Subventricular Zone (SVZ). The final aDEG set were those with an FDR-BH adjusted value less than 0.05. Graphics created with Biorender.com. **(b)** Distribution of 151,647 accepted nuclei recovered per condition, shown by age group and sex, demonstrates approximately even recovery across categories. **(c)** Nuclei distribution shown by brain region and age group (see also Supp. Table 2). (**d)** Clustering of 151,647 nuclei using the Leiden algorithm (see Methods) resulted in 25 unique clusters representing 10 unique broad cell-types. Clusters are colored and labeled according to broad cell-type annotation and **(e)** brain region of origin. Initial broad cell-type annotations were made using a combination of *Pegasus*-determined cluster markers (see Methods) and relative expression levels of known canonical marker genes for various CNS cell-types (Supp Fig. 2; Supp. Tables 3-4) This process revealed 3 neuronal populations (InN: inhibitory neurons, ExN: excitatory neurons, SPN: spiny projection neurons), 3 glial populations (Micro: microglia, Astro: astrocytes, Epend: ependymal cells), 2 oligodendrocyte-lineage populations (OPC: oligodendrocyte precursor cells, Oligo: oligodendrocytes), 1 endothelial (Endo), and 1 mural cell population each. **(f)** The proportions of ten unique cell-types found across the SVZ, PUT, MTG, and EC by age group. From the accepted 151, 647 nuclei, percentages were calculated using raw cell counts within a given region by age subset (n=5-7 individuals per region per age group). **(g)** Regional specificity of cell counts shows even distribution of cell-types across all regions of interest, except SPN which are known to be restricted to midbrain regions^23^. Regional specificity was calculated as a proportion of the number of a particular cell-type in the region of interest (ROI) over the total number of cells of that type (e.g. # ExN_MTG / # ExN_total). **(h)** Age effect on cell-type proportions reveals no effect (FDR > 0.05) in any cell-type across all 4 ROIs. Relative cell-type proportions were compared with a t-test and FDR-BH corrected to assess significance.

After quality control and pre-processing, we captured 151,647 nuclei, with similar recovery across age groups (n=75,547 young; 76,100 aged) and sexes (n=76,084 male; 75,563 female). Relative nuclei proportions by sex within each age group are shown in Figure 1B. By region, the nuclei distribution was 45,688 EC, 25,026 MTG, 34,444 SVZ, and 46,489 PUT. Relative nuclei proportions by age within each region are shown in Figure 1C. Raw nuclei counts by age for each cell-type and region are provided in Supplementary Table 2.

Using a combination of computed cluster marker genes and known CNS cell-type marker genes (Supp. Table 3-4), 25 distinct clusters were annotated into 10 broad cell-types (Supp. Fig. 2), including inhibitory neurons (InN), excitatory neurons (ExN), spiny projection neurons (SPN), oligodendrocyte precursor cells (OPC), oligodendrocytes (Oligo), astrocytes (Astro), microglia (Micro), ependymal cells (Epend), endothelial cells (Endo), and mural cells (Fig. 1D; see full cell taxonomy in Supp. Table 5). As expected, we found that some cell-types, such as microglia and oligodendrocyte precursor cells, were uniformly represented across all regions, while others, such as spiny projection neurons and excitatory neurons, showed greater regional specificity (Figure 1E). These observations indicate that we were able to generate a valid dataset to examine age-related changes in gene expression across multiple regions of the human brain.

### Cell-type proportions show no change across age groups

We investigated the effect of regional biases and age on cell-type proportions (Fig. 1F; see Methods for calculation). In total, 17,733 nuclei were annotated as inhibitory neurons (n=8,734 young; 8,999 aged), and we found that these nuclei predominate in the MTG (26.23% young; 22.94% aged) and EC (19.87% young; 19.89% aged; Fig. 1F). Similarly, excitatory neurons (n=11,757 young; 16,726 aged) were also recovered at high levels in the MTG (39.69% young; 54.81% aged) and EC (33.34% young; 37.17% aged; Fig 1F). A total of 29,967 cells were labeled as spiny projection neurons (17,473 young; 12,494 aged), and all were localized to the PUT (Fig. 1F). While there were initially nuclei annotated as spiny projection neurons outside of the PUT, we considered such assignments to be unreliable and likely due to small related subclusters of inhibitory neurons, to which SPN are a specialized subclass ^20^. Therefore, any non-PUT cells initially annotated as SPN were re-classified as “Other” (n=3,545 cells; Supp. Fig. 3) and excluded from further analyses. While the 50,129 cells (n=24,545 young; 25,584 aged) annotated as oligodendrocytes were present in all regions, the majority originated from the SVZ (75.35% young; 77.91% aged; Fig. 1F). These regional distinctions are further quantified via a regional specificity value (see Methods) that show that all other cell-types were uniformly distributed across regions (Figure 1G). We report no statistically significant effect of age to any cell-type proportion across all regions (FDR-BH adj. P-value >0.05 for all; Figure 1H; see Methods for proportion analysis), noting that sample size is relatively modest and does not allow us to fully exclude that there may be some changes in cell number. Based on these observations, we therefore tested the effect of age on gene expression within cell-type by region subgroups.

### Thousands of genes are differentially expressed across age groups

Gene sets were tested for age-related differential expression in a two-step process. We performed an initial t-test to identify strongest effect size candidates followed by a generalized linear mixed model with Tweedie distribution to more rigorously identify aDEGs (see Methods) followed by Benjamini-Hochberg False Discovery Rate (FDR-BH) correction for this more limited multiple testing. Across all cell-types, 8,872 unique genes were age-associated via linear model (Supp. Table 5). The majority of aDEGs were found to be protein coding, with the remaining being long non-coding RNAs (Supp. Fig. 4; Supp Table 6). While the number and direction of aDEGs are region- and cell-type-specific, the majority of aDEGs showed decreased expression with age (Supp. Fig. 5; Supp. Table 6). Based on these observations, we next examined how aDEGs were distributed in different cell-types.

### Inhibitory neurons demonstrate the greatest age effect among the neuronal cell-types across regions including dysregulation of protein synthesis

Within neuronal cell-types, inhibitory neurons were the most affected by age, with 2,708 aDEGs. Of these inhibitory neuron aDEGs, 2,444 were specific to a single region, with the majority being in the EC (n=1,216; Fig. 2A). In excitatory neurons, we identified 1,715 aDEGs. Of these, 1,613 were contained within a single region with the most being found in the EC (n=937; Fig. 2B). In PUT-specific spiny projection neurons, we found 768 aDEGs (Fig. 2C). All aDEGs for these cell-types can be found in Supplementary Table 6.

**Figure 2:**
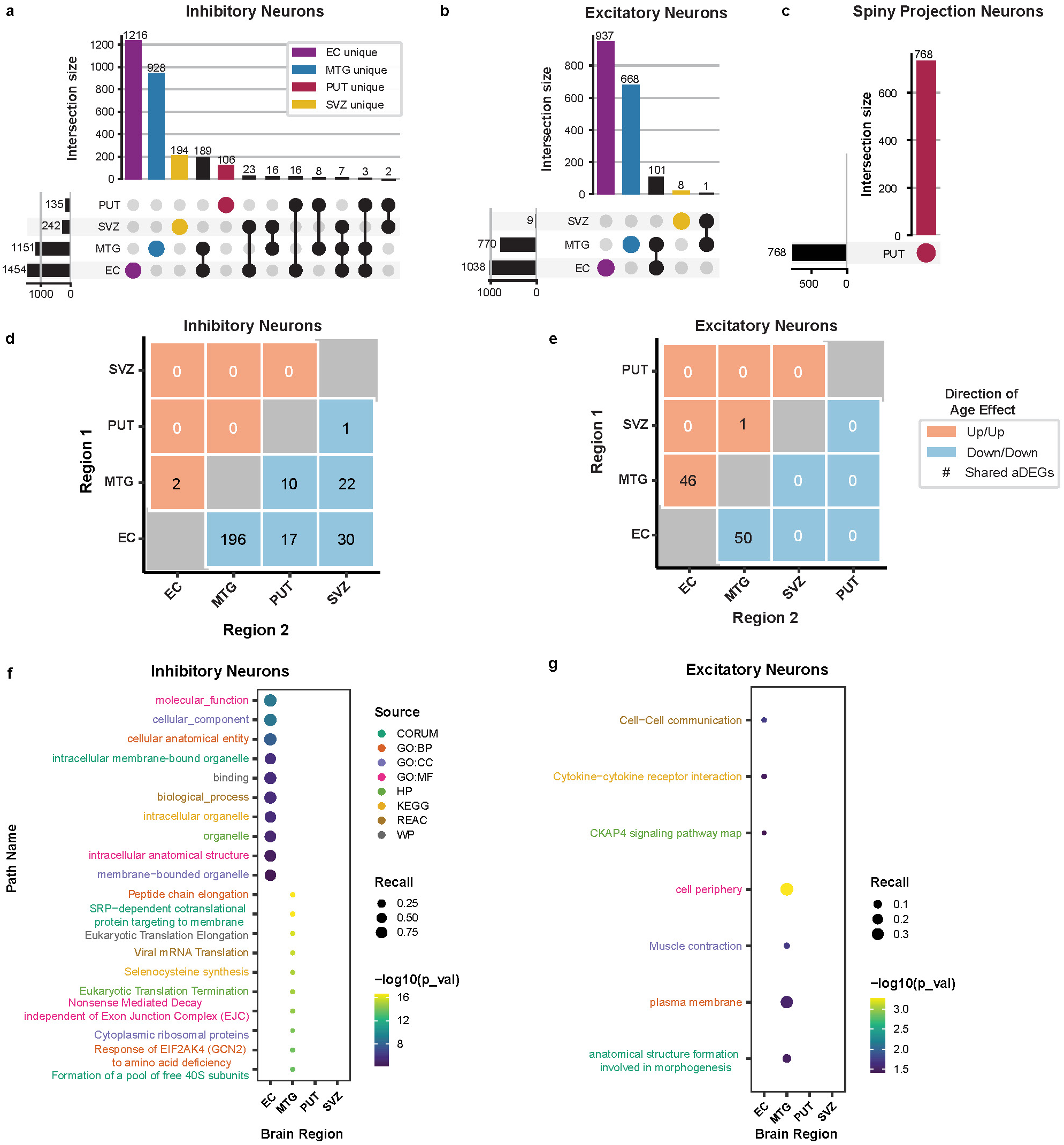
Inhibitory neurons exhibit the largest number of age-associated differentially expressed genes (aDEGs) amongst neuronal cell-types. Regional distribution and overlap of neuronal cell-type aDEGs indicate that the majority of aDEGs are unique to a particular region. **(a)** Inhibitory neurons yielded the largest number of aDEGs with the most being found in the EC. **(b)** Excitatory neurons had fewer total aDEGs but followed similar trends. **(c)** Spiny projection neurons are known to be biologically distinct to the putamen^23^, so they were only reported for this region (see Fig. 1H). Pairwise comparison of shared aDEGs within neuronal cell-types suggests similar signatures between the EC and MTG. Heatmap values representing count of aDEGs indicate the number of aDEGs changing in the same direction (concordance)--either increasing (positive, red) or decreasing (negative, blue)–in both of the indicated regions. **(d)** Inhibitory neurons showed 196 negatively concordant aDEGs between the EC and MTG. **(e)** Excitatory neurons showed 96 concordant aDEGs between the EC and MTG, with 46 positive and 50 negative. Top 10 significantly enriched pathways (FDR < 0.05) per brain region for neuronal cell-type aDEGs. Functional enrichments calculated in gProfiler with cell-type by region background correction (see Methods). **(f)** Inhibitory neurons exhibited functional enrichments for pathways involving cellular components and translation/protein synthesis pathways. **(g)** Excitatory neurons exhibited functional enrichments for pathways involving cellular components as well as cell communication and signaling.

The proportion of aDEGs that were shared across regions for neurons ranged from 1.1% to 11.4%. The majority of these shared-aDEG relationships occurred between two brain regions (pairwise). Generally, the direction of age-effect was found to be concordant in direction across regions and cell-types. In inhibitory neurons, the most striking of these relationships were 196 aDEGs shown to decrease in both the EC and MTG (negatively concordant; Fig. 2D). In contrast, for excitatory neurons, there were approximately even numbers of aDEGs that were increased (positively concordant; n=46) or decreased with age (negatively concordant; n=50) across both the EC and MTG (Fig. 2E). aDEGs that showed increased expression with age in one region but decreased expression in another (discordant relationships) were present but much fewer in number relative to concordant relationships (Supp. Fig. 6). In addition, while there were instances of aDEGs being shared across more than two regions, these were relatively infrequent.

There were 96 enriched pathways across all neuronal cell-types, with 88 observed in inhibitory neurons, all of which were restricted to the EC (n=20) or MTG (n=68; Supp. Table 7). Within the EC, the top inhibitory neuron pathways were related to cellular anatomy and subcellular components (i.e. organelles; Fig. 2F). Within the MTG, significant inhibitory neuron pathways related to translational processes, protein synthesis, and ribosomal structures (Fig. 2F). Excitatory neurons had seven total enriched pathways, all of which were also restricted to the EC (n=3) or MTG (n=4; Supp. Table 7). In the EC, we observed excitatory neuron pathways related to cell-cell communication, particularly relevant to cytokine receptor interactions and *CKAP4* signaling (Fig. 2G). In the MTG, excitatory neuron pathways related to cellular morphology–specifically the plasma membrane–and muscle contraction (Fig. 2G). PUT-specific spiny projection neurons were enriched for “*MECP2* and associated Rett syndrome” (WikiPathways:WP3584; Supp. Table 7). Taken together, these observations suggest distinct transcriptional changes across neuronal cell-types and regions with aging.

### Oligodendrocyte-lineage cells exhibit regionally distinct aging signatures related to neuronal structure and vesicular transport

In oligodendrocyte precursor cells (OPCs), there were 1,030 aDEGs (Supp. Table 6), with the majority of these aDEGs restricted to a single region (n= 913), particularly the EC (Fig. 3A). Mature oligodendrocytes yielded 2,406 aDEGs, 2,203 of which were specific to a single region with just under half of these (n=1,056) being specific to the MTG (Fig. 3B; Supp. Table 6). These findings suggest that oligodendrocyte precursor cells and mature oligodendrocytes show regionally distinct aging profiles. These distinct aging profiles are also observed when examining concordance in expression direction for aDEGs shared pairwise between regions. In OPCs, the aDEGs shared across regions were concordant in a negative direction, and the regions exhibiting the most aDEG overlap are the EC and the PUT (n=47), followed by the EC and SVZ (n=37; Fig. 3C). In contrast, for mature oligodendrocytes, there was a more even distribution between positively concordant and negatively concordant instances with the regions of greatest overlap being the EC and MTG (n=53, negatively concordant) and the SVZ and MTG (n=33; Fig. 3D). Discordant relationships can be found in Supplementary Figure 6.

**Figure 3:**
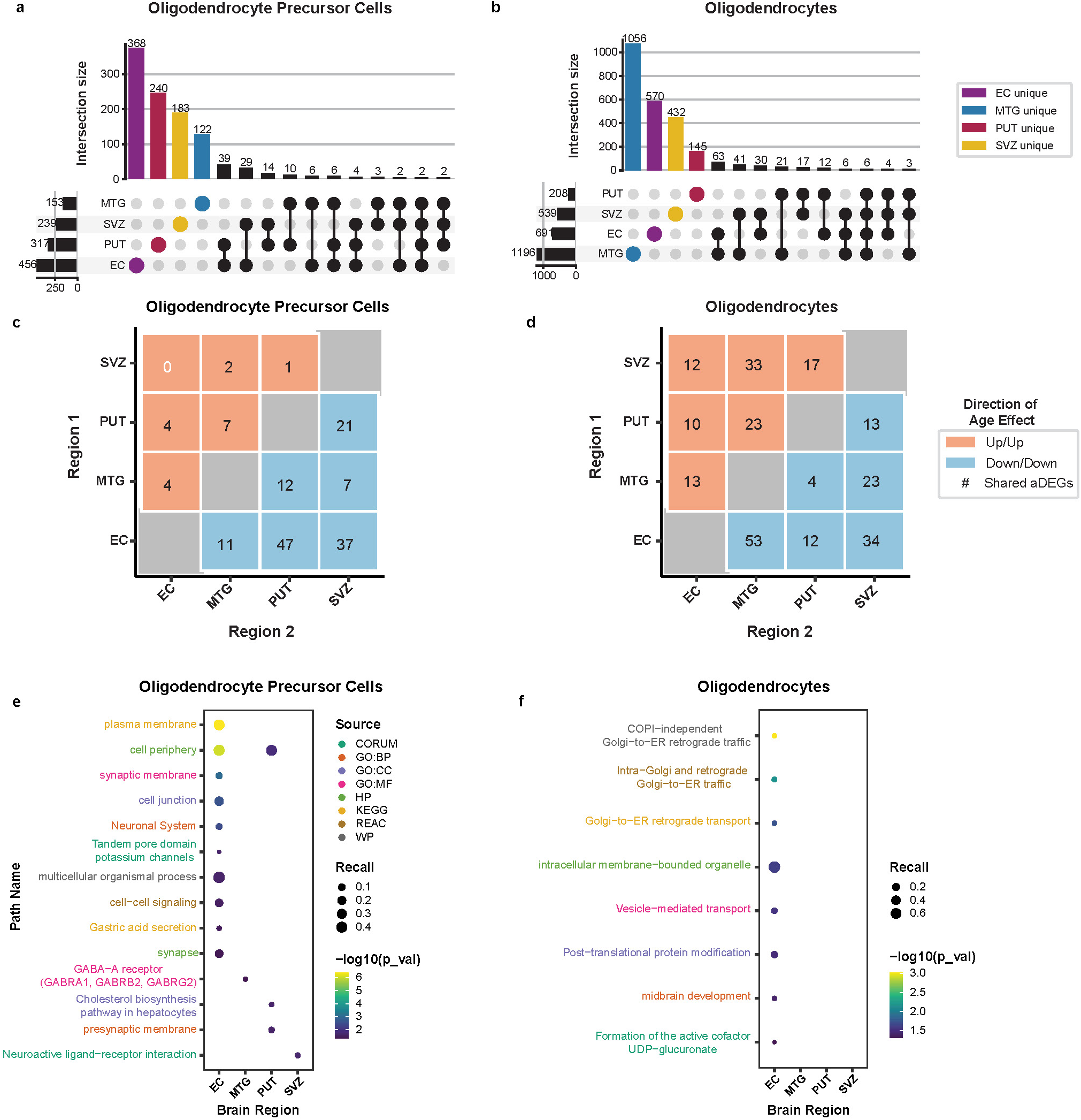
Oligodendrocyte-lineage cell aDEGs are enriched for neuronal cellular component & vesicular transport pathways. Regional distribution and overlap of neuronal cell-type aDEGs indicate that the majority of aDEGs are unique to a particular region. Of the oligodendrocyte-lineage cell-types, **(a)** oligodendrocyte precursor cells had its greatest proportion of aDEGs in the EC. **(b)** Mature oligodendrocytes had far more aDEGs, with the majority in the MTG. Pairwise comparison of shared aDEGs within oligodendrocyte lineage cells suggests variation between oligodendrocyte precursor cells and mature oligodendrocytes. Heatmap values representing count of aDEGs indicate the number of aDEGs changing in the same direction (concordance)--either increasing (positive, red) or decreasing (negative, blue)–in both of the indicated regions. **(c)** Oligodendrocyte precursor cells concordant aDEGs were largely in the negative direction. **(d)** Mature oligodendrocytes had a more balanced split between positively and negatively concordant aDEGs. Top 10 significantly enriched pathways (FDR < 0.05) per brain region for oligodendrocyte-lineage cell-type aDEGs. Functional enrichments calculated in gProfiler with cell-type by region background correction (see Methods). **(e)** OPCs showed enriched pathways for various neuronal cellular components and receptor pathways. **(f)** Mature oligodendrocytes showed enriched pathways related to vesicular transport.

Across oligodendrocyte-lineage cells, twenty-four enriched pathways were identified (n=16 OPC; n=8 Oligo; Supp. Table 7.). In oligodendrocyte precursor cells, significantly enriched pathways were found across all four regions, with the most being found in the EC (n=10; Fig. 3E; Supp. Table 7). Of these EC-OPC pathways, several relate to neuronal cell structures (e.g. plasma membrane, cell periphery, synaptic membrane, cell junction, neuronal system, potassium channels, and synapse). PUT-OPC also showed enriched pathways related to cell periphery and the pre-synaptic membrane. Oligodendrocyte precursor cells in the MTG and SVZ were enriched for the GABA-A receptor (CORUM:5809) and the neuroactive ligand-receptor interaction (KEGG:04080), respectively (Fig. 3E; Supp. Table 7). In mature oligodendrocytes, enriched pathways were related to Golgi-to-ER retrograde transport of vesicles and were restricted to the EC (Fig. 3F; Supp. Table 7). These results suggest that, in addition to neurons, substantial associations exist between aging and gene expression in oligodendrocytes and their precursor cells.

### Astrocytic and microglial cell aDEGs are found largely in the SVZ and EC and exhibit diverse functional enrichments

In astrocytes, a total of 1,717 genes were classified as age-associated (Supp. Table 6). Similar to other cell-types, the majority of these astrocyte aDEGs were unique to a single region, particularly in the EC (n=729; Fig. 4A). Microglia had a total of 676 age-associated genes, and over half were unique to the SVZ (n=344; Fig. 4B; Supp. Table 6). Ependymal cells had 479 aDEGs that were limited to the SVZ and EC, and aDEGs unique to the SVZ comprised most of these associations (n=374; Supp. Table 6).

**Figure 4:**
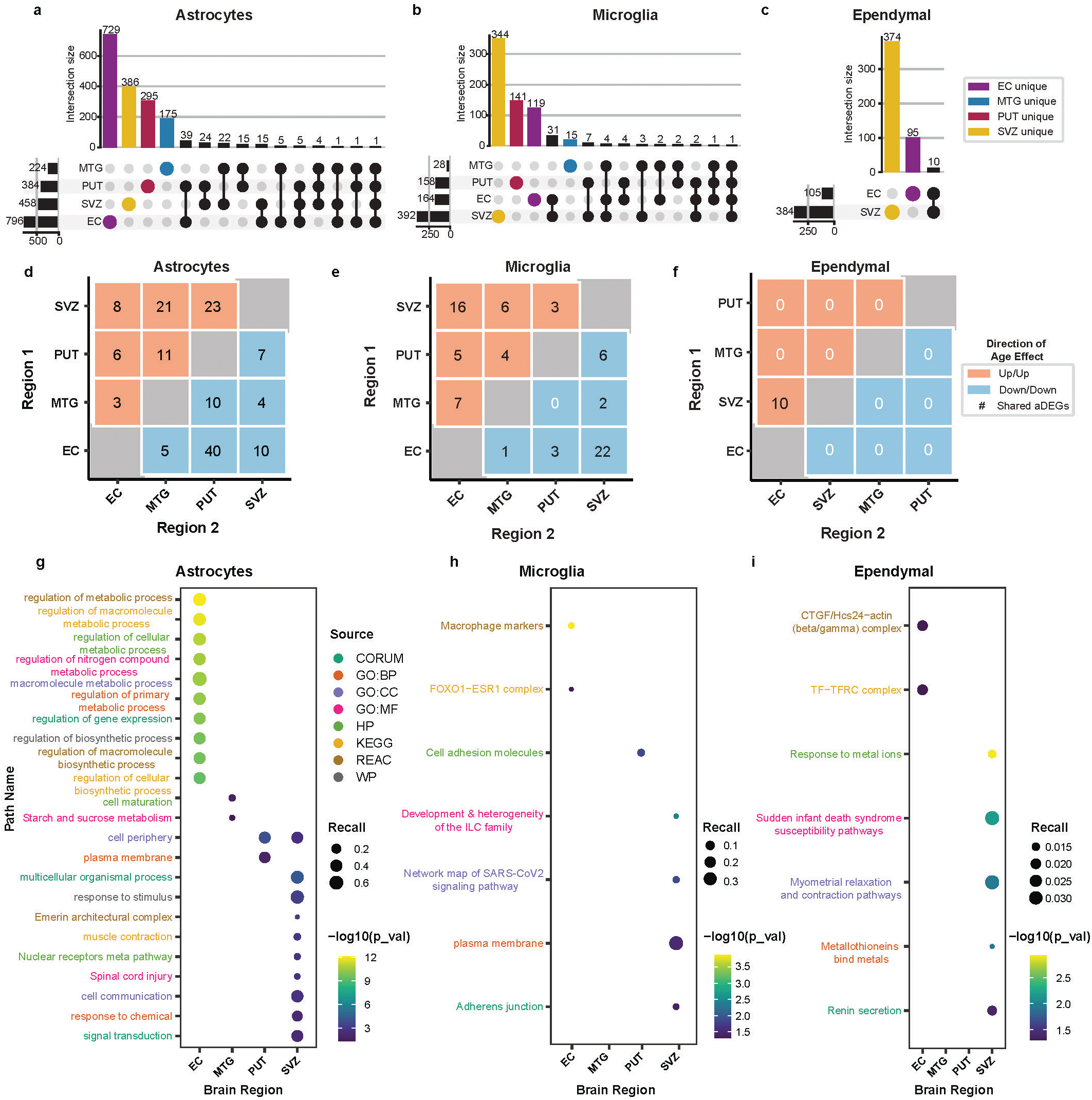
SVZ and EC account for the largest proportion of glial cell aDEGs and exhibit diverse functional enrichments. Regional distribution and overlap of neuronal cell-type aDEGs indicate that the majority of aDEGs are unique to a particular region. **(a)** Astrocytes yielded the most aDEGs of the glial cell-types with the greatest number being found in the EC. **(b)** Microglia aDEGs were largely found in the SVZ. **(c)** Ependymal cells had the fewest aDEGs of the glial cells and were found to be contained to the EC and SVZ. Pairwise comparison of shared aDEGs within glial cell-types suggests similar signatures between the EC and SVZ. Heatmap values representing count of aDEGs indicate the number of aDEGs changing in the same direction (concordance)--either increasing (positive, red) or decreasing (negative, blue)–in both of the indicated regions. **(d)** Astrocytes had a relatively large number of negatively concordant aDEGs across the EC and PUT. **(e)** Microglia showed their greatest numbers of concordant aDEGs between the SVZ and EC in both the positive and negative directions. **(f)** Ependymal cell concordant aDEGs were solely positive and contained to the SVZ and EC. Top 10 significantly enriched pathways (FDR < 0.05) per brain region for glial cell-type aDEGs. Functional enrichments calculated in gProfiler with cell-type by region background correction (see Methods). **(g)** Astrocytes showed enriched pathways related to metabolism, gene expression, biosynthesis, and signaling. **(h)** Microglia showed enriched pathways related to macrophage markers, immune cell development, and cell adhesion. **(i)** Ependymal cells showed enriched pathways relating to various cellular complexes and binding or response to metal ions.

Considering the concordance in direction between shared aDEGs, each glial cell population exhibits unique regional sharing profiles. Among astrocytes, there was a split between positive and negative instances of concordance (Fig. 4D). Notably, there were 40 negatively concordant aDEGs shared between the EC and PUT and the positively concordant aDEGs shared between the SVZ and PUT (n=21) as well as the SVZ and MTG (n=23; Fig. 4D). In contrast, for microglia, the greatest amount of overlap was between the SVZ and EC for both positively (n=16) and negatively concordant (n=22) aDEGs (Fig. 4E). For ependymal cells, all 10 shared aDEGs were positively concordant and were found in both the SVZ and EC (Fig. 4F). Discordant relationships can be found in Supplementary Figure 6.

Glial cell aDEG sets yielded 192 enriched pathways with the vast majority being in astrocytes (n=178), particularly within the EC (n=151; Supp. Table 7). In the EC astrocytes, the top enriched pathways were linked to metabolic processes, biosynthesis, and regulation of gene expression (Fig. 4G). In the SVZ, astrocyte pathways were more diverse and were linked to cell-cell communication, signal transduction, and response to stimuli (Fig. 4G). PUT astrocyte pathways appear to be specific to the plasma membrane, while MTG astrocyte pathways are involved in cellular maturation and starch/sucrose metabolism (Fig. 4G). In microglia, there were seven total enriched pathways across all regions (Supp. Table 7). The most significantly enriched pathway was for macrophage markers in the EC (Fig. 4H; Supp. Table 7). A connection to the immune system was also seen in SVZ microglia, which were enriched for Innate Lymphoid Cell (ILC) family development (WikiPathways:WP3893; Fig. 4H). Cell-cell adhesion was also found as a theme in SVZ and PUT microglia enriched pathways (Fig. 4H). Ependymal cells had a total of seven enriched pathways with the most being in the SVZ (Fig. 4I; Supp. Table 7), of which the most significant was response to metal ions, while another pathway implicated metallothioneins (Fig. 4I; Supp. Table 7).

Endothelial and mural cells exhibited relatively few aDEGs (n=47 and n=89, respectively; Supp. Fig. 7A-B; Supp. Table 6), with over half specific to the entorhinal cortex. Shared aDEGs were rare and limited to pairs of regions (Supp. Fig. 7C-D), with no discordant or broadly shared genes.

## Discussion

Transcriptomic analysis of postmortem brain samples is integral to the study of age-related diseases; however, little is known about how cell-types change during normal aging. Although a limited number of age-matched controls are available across several studies^5,6,25–28^, these are largely collected from the prefrontal cortex. Importantly, it is unknown whether cell-types respond to non-pathological aging similarly across regions. Therefore, there is a need for studies that intentionally target the aging process across multiple brain regions within the same donors. To address these gaps in literature, we performed single-nucleus RNA sequencing in cohorts of young (20-30 years old) and aged (60-85 years old) individuals across four regions–the entorhinal cortex (EC), middle temporal gyrus (MTG), subventricular zone (SVZ), and putamen (PUT).

Prior studies on aging using bulk RNA sequencing reported a decrease in cellular marker gene expression^3,24,29^, but a decrease in gene expression may not necessarily correspond to a physical loss of cells^24,30^. Here, we did not find that age had a significant effect on cell-type proportions across any broad cell-type or region (Fig. 1). Although there is some disagreement in the field regarding whether sampling via single-cell RNA sequencing is an appropriate method for cell counting^31,32^, other studies have large cell proportion shifts with age in rodents^33–37^, in animal models of disease^38–45^, and in postmortem human tissues across disease conditions^46–50^. We found relatively limited changes in cell numbers. This discrepancy may arise because studies tend to make conclusions about cell loss from data reporting a reduction in cortical volume, which could be suggestive of a reduction in myelination or cell size rather than an overt loss of cells^22,51^. Moreover, stereological counts of specific cell-types are often done in the context of highly-specialized regions, and these studies have shown variability based on the species, the chosen brain region(s), and the cell-types investigated^21,23,24,35,52–56^. However, we note that we have a relatively small sample size and that larger studies are needed to draw firm conclusions.

Prior reviews of transcriptional changes with age have emphasized that decreases in synaptic gene expression and increases in microglial activation were common, particularly in the pre-frontal cortex^2^. Consistent with these findings, we have previously reported a decrease in gene expression modules associated with synaptic genes and an increase in modules associated with inflammation^3^. However, most of these claims derive from microarray or bulk RNA-seq data, both of which lack single-cell resolution. By compiling a single cell dataset from multiple regions, we describe differentially expressed genes and pathways dysregulated across age on the broad cell-type level for each region. Overall, we report that age-associated differentially expressed genes (aDEGs) tend to be unique to a particular brain region. While we found that aDEGs decrease in inhibitory neurons, inflammatory signals, and other aDEGs tend to be region- and cell-type-specific without much overlap. The cell-type-specific (e.g. microglia compared to astrocytes) as well as region-specific (e.g. SVZ microglia compared to EC microglia) effects reported in our study highlight the importance for atlas-level, multi-region profiling of non-pathological aging.

Our study profiled three broad neuronal cell-types, consisting of inhibitory (InN), excitatory (ExN), and spiny projection neurons (SPN; Fig. 2). We found that inhibitory neurons exhibited the most aDEGs of any neurons with the largest proportion being in the EC. Of the genes that were shared across regions in inhibitory neurons, the most notable were the 196 aDEGs that decreased in expression with age in both the EC and MTG. Inhibitory neuron aDEGs were functionally enriched for translation and biosynthesis of proteins, particularly in the MTG. Many of the intersecting genes within these pathways encode ribosomal proteins (Supp. Table 7). Speculatively, it is possible that impaired ribosomal function leads to dysregulation of translation, potentially resulting in abnormal protein synthesis. These findings align with those of previous studies that have implicated aberrant proteostasis and ribosomal dysfunction in the aging process^18,57–61^. Our study presents the added layer of attributing this dysfunction specifically to inhibitory neurons, which could result in downstream impairments to inhibitory signaling and subsequently disrupt excitatory-inhibitory balance^62–66^. Excitatory neurons showed similar trends to the inhibitory neurons in that most of their aDEGs were specific to the EC, and that the regions with the most pronounced aDEG sharing were the EC and MTG. However, excitatory neurons differed in that they also showed a large number of aDEGs with an age-dependent increase in expression. Given that we observed region-specific age effects are present even in cell-types that are not altered in proportion, our findings lay a groundwork for understanding how different neurons of the brain respond differently to age.

For oligodendrocyte-lineage cells, we found that the aging profiles differed between oligodendrocyte precursors and mature oligodendrocytes (Fig. 3). For example, the MTG had the most region-specific aDEGs for mature oligodendrocytes but the fewest region-specific aDEGs in oligodendrocyte precursor cells. Moreover, both subtypes of oligodendrocyte-lineage cells also differed in their functional enrichments in that oligodendrocyte precursor cell aDEGs showed enriched pathways involving neuronal cell components (e.g. membranes, synapse, receptors) while mature oligodendrocyte enriched pathways highlighted post-translational modification, vesicular trafficking/transport, and development. The enrichment of neuronal cell components in the oligodendrocyte precursor cells aDEG sets could point towards a dysregulation of OPC-neuronal interactions in aging. OPC-mediated myelination and OPC-neuron synapses have both been shown to decrease with age^67–69^, which could have implications for modulation of neural circuit activity^69,70^. The oligodendrocyte enriched pathways for vesicular trafficking may also relate to neuronal myelination because this process requires proper lipid delivery mediated by SNARE complex VAMPs^71–73^. In addition, vesicular trafficking has also been shown to regulate oligodendrocyte maturation post-differentiation^74^. Given that our study reported overrepresentation of trafficking processes, this may suggest that oligodendrocyte maturation and myelination are altered during aging, which aligns with previously reported age-related decreases in white matter volume^21,75–77^.

The other glial cell-types represented in our study were astrocytes, microglia, and ependymal cells, and each showed a diverse aging profile (Fig. 4). Astrocytes had the most aDEGs of this subgroup and showed the most significantly enriched pathways of any cell-type, with the largest effect in the EC. These pathways showed clear trends for biosynthesis, gene expression, and several metabolic pathways. Astrocytes use glycolysis for generation of lactate as an efficient energy source to support the high metabolic demands of neurons ^78,79^. This metabolic profile has been reported to shift as a result of aging^80–83^, which could be reflected by changes to relevant gene expression. In contrast to astrocytes, both microglia and ependymal cells showed the most region-specific aDEGs in the SVZ. Of note, these ependymal cell aDEGs were functionally enriched for pathways related to the binding of metal ions. This is consistent with both an emerging role of ependymal cells^84–86^ and that this homeostatic process is disrupted during typical aging and in age-related diseases^87–90^.

Notably, our study is only the third single-nuclei study of the adult SVZ. Previous studies focused either on aged-individuals alone^91^ or the comparison between young and middle aged^92^. Our report that there are large age-related shifts in glia is consistent with these two previous studies, however the region remains understudied despite its significant role in development^93,94^ and disease^95–100^. This is especially critical as the SVZ would be the site of adult neurogenesis^16,101–103^, and there is evidence suggesting that impaired neural stem cell function may be implicated in neurodegenerative disorders^104–106^. The largest of the two previous studies compared age groups of 16-22 (young) and 44-53 (middle-aged) with particular attention to developmental processes^92^. They found that while neural stem cell numbers did not decline across their sample ages, oligodendrocyte precursor cells and microglia did, along with an associated decrease in developmental genes. The authors suggested that these findings pointed to a remodeling of the SVZ between youth and adulthood. While our study is similarly powered in terms of donors and cells captured, the disparity in findings may relate to the specific chosen age range in the older group (Puvogel et al.: 44–53 years, current study: 60–85 years). As more SVZ samples become available and single-nuclei data are collected, future studies will be necessary to separate periods of development from any eventual decline with age.

Our sample series prioritizes large effect sizes across young (20–30 years old) and aged (60–85 years old) cohorts and represents the largest multi-region, non-pathological brain aging study to date (n=5–7 donors x 4 regions). One limitation of this design is the lack of middle-aged samples, limiting our ability to appreciate gene expression changes that may have complex, continuous relationships with chronological age. Furthermore, technologies to sample gene expression are the most routinely used methods to profile unbiased genome-wide changes at single cell resolution. As single cell and spatial technologies expand to include other readouts such as global proteomics, DNA accessibility, and nucleotide modifications, these additional multi-omic modalities have the potential to enhance our understanding of how gene-regulation changes with age.

## Data and Code Availability

Raw single-nucleus RNA sequencing data are available in the NIMH Data Archive associated with the collection “Human Brain Collection Core genomics data in postmortem brain of psychiatric disorders #3151” (https://nda.nih.gov/edit_collection.html?id=3151) experiment ID 2370: “snRNA_brain_aging.” De-identified individual level meta-data used as either selection criteria and/or covariates in analysis can be found in Supplementary Table 1. Summary-level results used to create figures are available on Zenodo (https://zenodo.org/records/15800429) and in Supplementary tables 7-9. Associated code used for analysis are available on GitHub (https://github.com/neurogenetics/ADRD_Brain_Aging/tree/main/phase1).

## Acknowledgments

The tissue / data used in this research was obtained from the Human Brain Collection Core, Intramural Research Program, NIMH (http://www.nimh.nih.gov/hbcc) and sequencing was performed by the NIH Intramural Sequencing Core (NISC). This research was supported by the Intramural Research Program of the National Institutes of Health, National Institute on Aging project 1ZIAAG000539-01 (to M.R.C), and in part by project ZO1 AG000535 (to M.A.N.); as well as the National Institute of Neurological Disorders and Stroke (1ZIANS003154, S.W.S.). The HBCC is supported by NIMH_IRP project # ZIC MH002903. This work utilized the computational resources of the NIH HPC Biowulf cluster (http://hpc.nih.gov).

## Author Contributions

M.E.M., M.D., R.L., S.W.S., J.R.G., and M.R.C., conceptualized the study and planned experiments. Investigation and formal analysis was performed by M.E.M., M.D., D.J.A., J.D., S.I.S., X.R., D.T.W., P.K.A., and J.R.G.. Funding of this study was obtained by M.R.C., and M.A.N.. M.E.M. and M.D. wrote the original draft. M.R.C., A.D., and S.M., supervised the work. All authors reviewed and edited the manuscript.

## Methods

### Brain Bank Information & Sample Selection

Brain tissue samples were obtained from the National Institute of Mental Health (NIMH)’s Human Brain Collection Core (HBCC). Brain donors were identified through the Office of the Chief Medical Examiner (OCME) of the District of Columbia and Northern District in Virginia, since the present study intended to investigate the molecular mechanisms of normal aging, we solicited tissue samples from individuals who had *not* been diagnosed with any neuropsychiatric disorder in their lifetime. In lieu of quantified neuropathology scores (i.e. Braak Tau, Braak Syn, Thal), selected samples did not have apparent evidence for neurodegeneration. Pathological assessment was performed by a neuropathologist. Procedures for assessment and characterization of cases are described in previous publications^107^.

Suitable donors had usable tissue across the four brain regions of interest–the entorhinal cortex (EC), middle temporal gyrus (MTG), subventricular zone (SVZ), and putamen (PUT). We selected individuals with the lowest available postmortem interval (PMI range: 13-58 hrs) and brain pH levels (range: 6.19-6.92). Samples were classified by age with younger being 20-30 years old (n=6; 3 per sex) and older being 60-85 years old (n=7; 4 male, 3 female) at time of death. Due to tissue availability or sample processing failure, three donors are not full-rank. However, each region has contributions from 12 independent donors (5-7 per age condition; see Supp. Fig. 1 and Supp. Table 1 for sample characteristics and demographic info.).

### Isolation of nuclei from human brain

Nuclei were isolated using the Nuclei PURE Prep Nuclei Isolation Kit (Sigma #NUC201) per the manufacturer’s instructions. To streamline the workflow and minimize potential batch effects, we utilized a pooling approach prior to library preparation and proceeded with post-sequencing demultiplexing, generating six pools of eight samples each. Each pool was balanced for age group and sex ensuring that no single pool contained two regions from the same donor.

A 100 mg piece of tissue was placed on a fresh, pre-chilled petri dish, trimmed, and weighed. Within each of the six pools, approximately 25 mg of tissue for each of the eight samples was combined and homogenized in a single cold douncer containing 2 mL ice-cold lysis buffer (Nuclei PURE Lysis Buffer (Sigma #L9286), 0.1M freshly-thawed DTT (Sigma #GE17-1318-01), and 0.1% Triton X-100 (Thermofisher Scientific #T1565). Tissue was homogenized with 25 strokes of a loose pestle followed by 25 strokes of a tight pestle, transferred to a tube containing 8mL cold lysis buffer, vortexed 2-3s, and left on ice for 10 minutes. Following lysis, cold 1.8M sucrose cushion solution (Nuclei PURE 2M Sucrose Cushion Solution (Sigma #S9308), Nuclei PURE Sucrose Cushion Buffer (Sigma #S9058) and 0.1M DTT) was added to the bottom of an ultracentrifuge tube (Beckman Coulter #344058) on ice. To each lysate, cold 1.8M sucrose cushion solution was added and mixed using a serological pipette. The lysate solution was slowly layered on top of the sucrose cushion, placed in a pre-cooled ultracentrifuge, then centrifuged for 45 min. at 30,000 x g at 4℃.

Sample tubes were removed from the ultracentrifuge and placed on ice. The entirety of the supernatant was aspirated, and the nuclei pellet was resuspended in Nuclei Suspension Buffer on ice (NSB; 1mL cold PBS (Thermofisher Scientific #10010-023), 0.01% BSA (New England Biolabs #B9000S), 0.1% SUPERase RNase inhibitor (Thermofisher Scientific #AM2696), transferred to a 15mL tube containing an additional 4mL NSB buffer, mixed, and washed by centrifugation at 500 x g for 5 min. at 4℃. The pellet was resuspended in 1 mL NSB, filtered through a 70μM Cell Strainer (STEMCELL Technologies #27216) to remove debris, and washed again via centrifugation. The supernatant was aspirated, and nuclei were resuspended in a final volume of 110μL.

To determine nuclei concentration, Acridine Orange/Propidium Iodide (Logos Biosystems #F23001) was added to nuclei suspension in a separate tube and counted using a LUNA-FL Dual Fluorescence Cell Counter (Logos Biosystems). An appropriate volume of nuclei was diluted with NSB to achieve a final single suspension of 80,000 nuclei (per sample) to maximize nuclei recovery while minimizing the multiplet rate. Approximately 10,000 nuclei from this single suspension were loaded into each of 8 lanes of a 10x Genomics NextGEM Chip G (10x Genomics #1000127) and inserted into a 10xGenomics Chromium Controller (10xGenomics #1000204) according to the manufacturer’s instructions. We loaded 10K nuclei per lane, targeting ∼6K nuclei recovered per lane with a multiplet rate of ≤5%. The remainder of library preparation was conducted according to the Chromium Single Cell 3’ Reagent Kits User Guide (v3.1 Chemistry; Rev.D). Final libraries were sequenced at the NIH Intramural Sequencing Center (NISC) at a read depth of 25,000 paired-end reads per nucleus on an Illumina NovaSeq 6000. Data was processed with CellRanger count v5.0.1 with refdata-gex-GRCh38-2020-A as the reference, with introns included.

### Genotype Preparation and Demultiplexing of Pooled Samples

#### Genotypes were assayed from three array-based genotyping platforms

Human1M-Duov3_B, HumanHap650Yv3.0, and HumanOmni5-Quad. Genotypes were merged and filtered to variants that were common to all three platforms, called in 95% of the samples, and known to be single-nucleotide variants (SNVs). Genotype files were lifted over from hg19 to hg38. The Plink2.0^108^, bcftools^109^(version 1.12), and Picard (version 2.25.5) (https://broadinstitute.github.io/picard/) tool sets were used to filter, format, and liftover the genotype files. Pooled samples were demultiplexed using the demuxlet tool (version 2)^110^. The prepared subject genotypes and the aligned single-nuclei bam files were used to deconvolute the cells’ sample identities. Data were processed on the Google Cloud Platform (GCP) using the Cumulus/Demuxlet workflow (WDL, https://cumulus-doc.readthedocs.io/en/0.12.0/demuxlet.html), which executes the demuxlet tool contained in the Statgen Popscle suite (https://github.com/statgen/popscle). Job submission to GCP for execution was done via the Broad WDL runner (https://github.com/broadinstitute/wdl-runner) and GCP Life Sciences interface (https://cloud.google.com/life-sciences/docs). SCANPY^111^ (version 1.7.1) was used to read in the 10X filtered matrix files into an AnnData object and integrate sample identity for the deconvoluted cells. Cells that were scored as ambiguous or doublets by genotype were excluded.

### Clustering and cell-type identification

The single-cell analysis tool Pegasus^19^ (version 1.3) (https://pegasus.readthedocs.io/en/stable/index.html) was used to combine, filter, normalize, cluster, and determine the initial cell-type identities of the demultiplexed single-nuclei count data. Basic filtering was done with Pegasus excluding cells that did not include at least 200 genes, genes that were not present in at least three cells, and cells that had a mitochondrial DNA content of over 10%. Clustering was performed on log-normalized counts using the top 2,000 variable genes. Harmony-corrected principal components were calculated and clustered using fast approximate nearest neighbor search (hnswlib^112^). The Leiden^113^ algorithm (version 0.8.3) was used to identify clusters on the neighborhood graph, and multiple resolutions were inspected to determine appropriate cell-type identification.

Initial inferences for the putative cell-type identity of each cluster were made based on a combination of the computed marker genes for that cluster compared to all others and the relative expression levels of canonical marker genes for different CNS broad cell-types (For a list of the marker genes, calculated with default settings in Pegasus, see Supp.Tables 3-4, Supp. Fig. 2). To preserve the number of expected cell-types in a single round of clustering, we selected a final Leiden resolution of 0.85. Cellbender^114^ was run to determine the degree to which ambient RNA may impact the cell data and clusters. Of the cells detected as ambient by Cellbender, 98.9% of these cells were filtered out by either CellRanger or demuxlet. For those filtered by demuxlet the majority were labeled as ambiguous. Of the small fraction of remaining ambient cells 76.6% of these were assigned to a cluster within the ‘Other’ cell-type; where 21.3% of cells within the ‘Other’ clusters were scored as ambient. In cases where a broad-cell-type annotation could not be definitively assigned via algorithmic or manual inspection or were determined to be biologically inappropriate (e.g. cells labeled as “Spiny Projection Neurons” but were localized to regions other than the Putamen), these cells were labeled as “Other” and excluded from further analysis (n= 16,298 nuclei). As a separate analysis (Supp. Fig. 3), sub-clustering of SPNs performed in Seurat^115^ and integrated with Harmony^112^ further demonstrates that dopaminergic neurons in this study are largely contributed by the putamen. The final accepted dataset consisted of 151,647 nuclei, clustered into 10 annotated cell-types.

### Cell-Type Proportion and Regional Specificity Analyses

In order to compare cell type proportions across age, nuclei counts by region per age group were obtained and this value was used to calculate the percentage of nuclei that this cell type comprised within that particular region and age group (see values in Supplementary Table 2; percentages in Figure 1F). To evaluate whether these differences in nuclei numbers between age groups were due to a significant effect of age on cell-type proportions, the *propeller*^116^ function of the R-Package *speckle* (version 1.60) was used with default parameters as recommended by the authors. In brief, this function applies a logit transformation to per-sample cell counts followed by a t-test to compare the age effect (young vs aged) in each cell-type. This method was applied within each region and adjusted the p-values using the Benjamini and Hochberg False Discovery Rate (FDR-BH) method prior to determining significance for all comparisons across the 4 regions. For ease of interpretation, summary proportions per cluster were then normalized (−1: 100% young; 1: 100% aged).

To assess whether any cell-types were more specific to a particular region, a regional specificity value was calculated for each cell-type by region combination. These specificity values were calculated by determining the total number of cells of a given type within the region of interest and dividing that by the total number of cells of that type across all regions (e.g. # ExN_MTG / # ExN_total). For example, if a cell-type-region combination had a specificity value of 0.5, 50% of all cells of that type were contained within that one region. Source code for determination and heatmap visualization of both proportion and regional specificity values can be found on GitHub.

### Age-related differential expression and association

Differential expression analysis was performed with age group as the independent variable per broad cell-type per region. Age was treated as a binary, categorical variable given as “young” (20-30 yrs.) or “aged” (60-85 yrs.). To exclude poorly mapped genes, we applied a filter requiring that genes tested for differential expression must have at least one read in a minimum of 3 cells for at least 50% of subjects. For computational efficiency, the differential expression analysis was performed in two steps.

First, a simple t-test between age groups was performed using the *diffxpy* package (https://diffxpy.readthedocs.io/en/latest/index.html). Any result exhibiting a nominally significant difference was considered to proceed to the second step. In this second step, to address impacts of pseudoreplication and zero-inflation, we utilized a generalized linear mixed model (GLMM) with a Tweedie distribution^117^. Specifically, to account for pseudoreplication, a random-effect term was included to account for the sample, while a Tweedie distribution was specified to account for zero-inflation. Additionally, the pool number was included as a term to account for residual batch effects that were not corrected with Harmony. The glmmTMB^118^ R package was used to run this model; gene ∼ age_group + pool + (1|sample_id). To correct for multiple testing across both cell-type and region, the resulting p-values were adjusted using the Benjamini and Hochberg False Discovery Rate (FDR-BH) method, as implemented in the statsmodel multitest Python package (https://www.statsmodels.org). In order to evaluate genes that were differentially-associated with age (aDEGs) in a particular cell-type, the Python package *UpSetPlot* (version 0.9.0) (https://upsetplot.readthedocs.io/en/stable/#) was used to visualize aDEG distribution and sharing across regions. All UpSet Plots are sorted by cardinality with no cut off parameters. The input dataframes for each UpsetPlot can be found in Supplementary Table 8.

### Evaluating Concordance/Discordance of aDEG Expression Direction across Regions

Given that there were aDEGs being shared across two or more brain regions within a given cell-type, we investigated whether the expression of these genes changed in the same or opposing directions with age between 2 regions of interest. A concordant signature had the aDEG increasing (up/up) or decreasing (down/down) with age in both regions. A discordant signature had the aDEG increasing in one region and decreasing in another (up/down or down/up). The input dataframes for each heatmap can be found in Supplementary Table 9.

### Functional Enrichment Analysis with Background Correction

Functional enrichment analysis was performed on each of the aDEG sets determined by the custom generalized linear mixed model (FDR BH p< 0.05) within each broad cell-type, brain region combination using the R package *gprofiler2*^119(^version 0.2.3). For each cell-type by region aDEG set, all genes were evaluated together, independent of effect direction (i.e genes increasing in expression with age were not evaluated separately from those decreasing) in order to preserve biological context. In addition, all reference gene sets for tested pathways were background-corrected for genes that were generally expressed (i.e. not an aDEG) in a given cell-type by region subset (i.e. they were removed from the reference set).

---

**Supplemental Figure 1:**
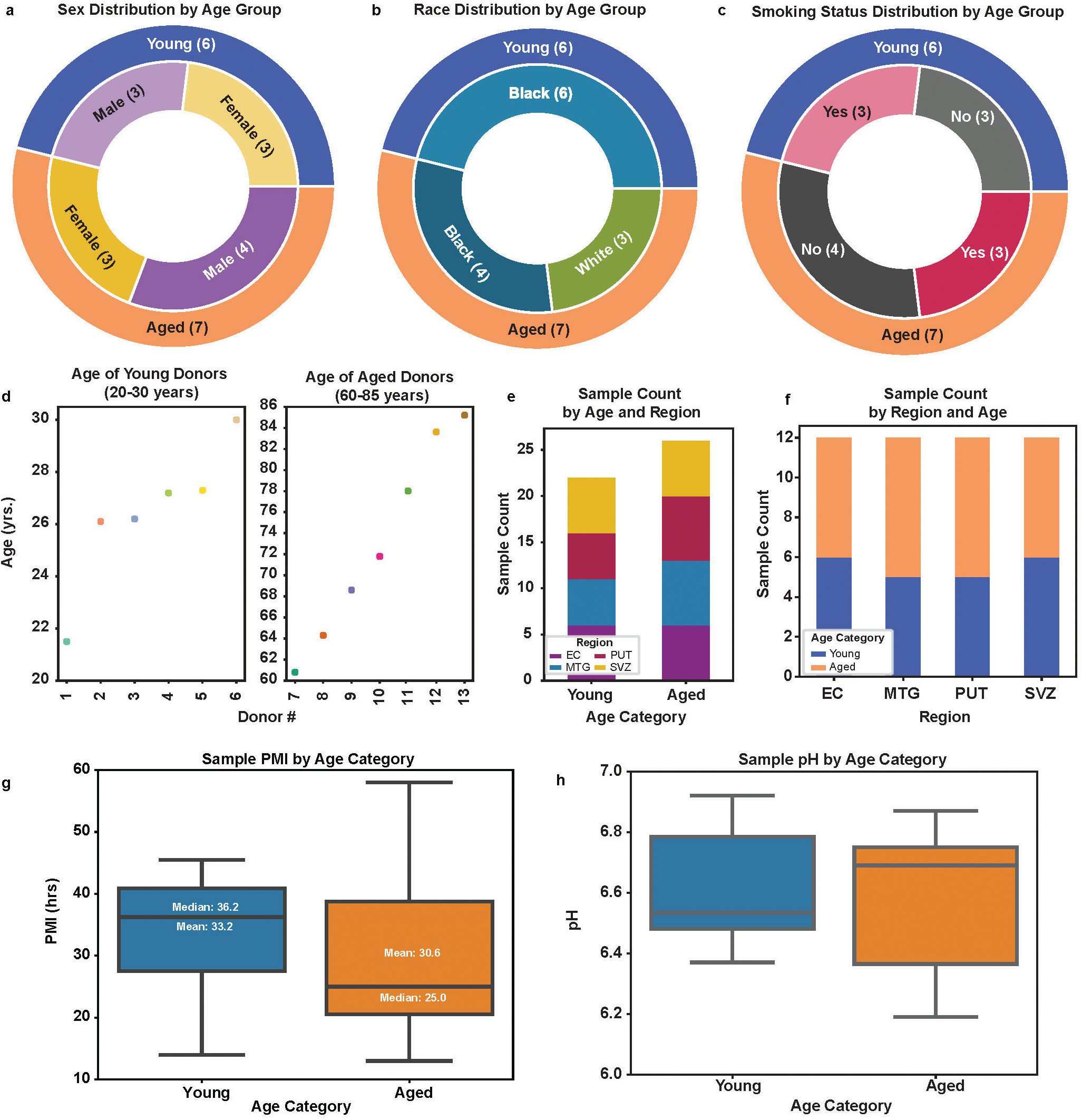
Donor Demographics and Sample Characteristics. Distribution of demographic characteristics across 13 donors within young (20-30 yrs.) versus aged (60-85 yrs.) groups: **(a)** sex, **(b)** race, **(c)** smoking status, and **(d)** age. Distribution of tissue sample counts **(e)** by region within an age category and **(f)** by age category within a region of interest demonstrates a relatively even sample distribution. This sample series was selected based on minimizing variability in **(g)** post-mortem interval (PMI; range 13-58 hrs; mean: 33.2 hrs.; median: 36.2 hrs.) and **(h)** brain pH (range 6.19-6.92).

**Supplemental Figure 2:**
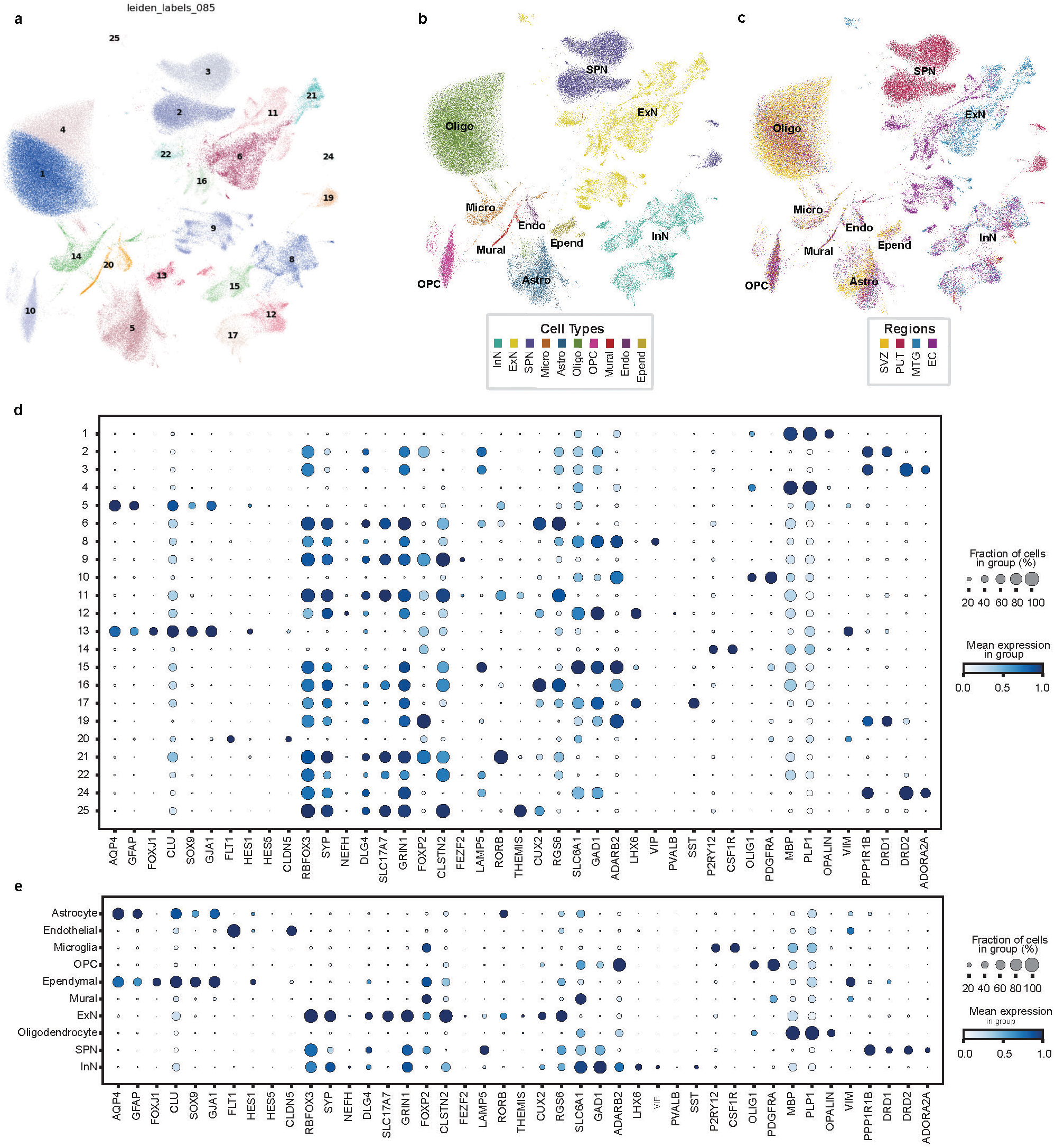
CNS Marker Gene Expression across 25 Leiden Clusters reveals 10 unique cell-types across four brain regions. **(a)** Leiden clustering of 151,647 nuclei at 0.85 resolution resulted in 25 distinct clusters. **(b)** Leiden clusters re-colored and labeled according to broad cell-type annotation and **(c)** brain region of origin. **(d-e)** Relative expression levels and proportion of cell-types expressing canonical marker genes were used to manually annotate to the broad cell-type level.

**Supplemental Figure 3:**
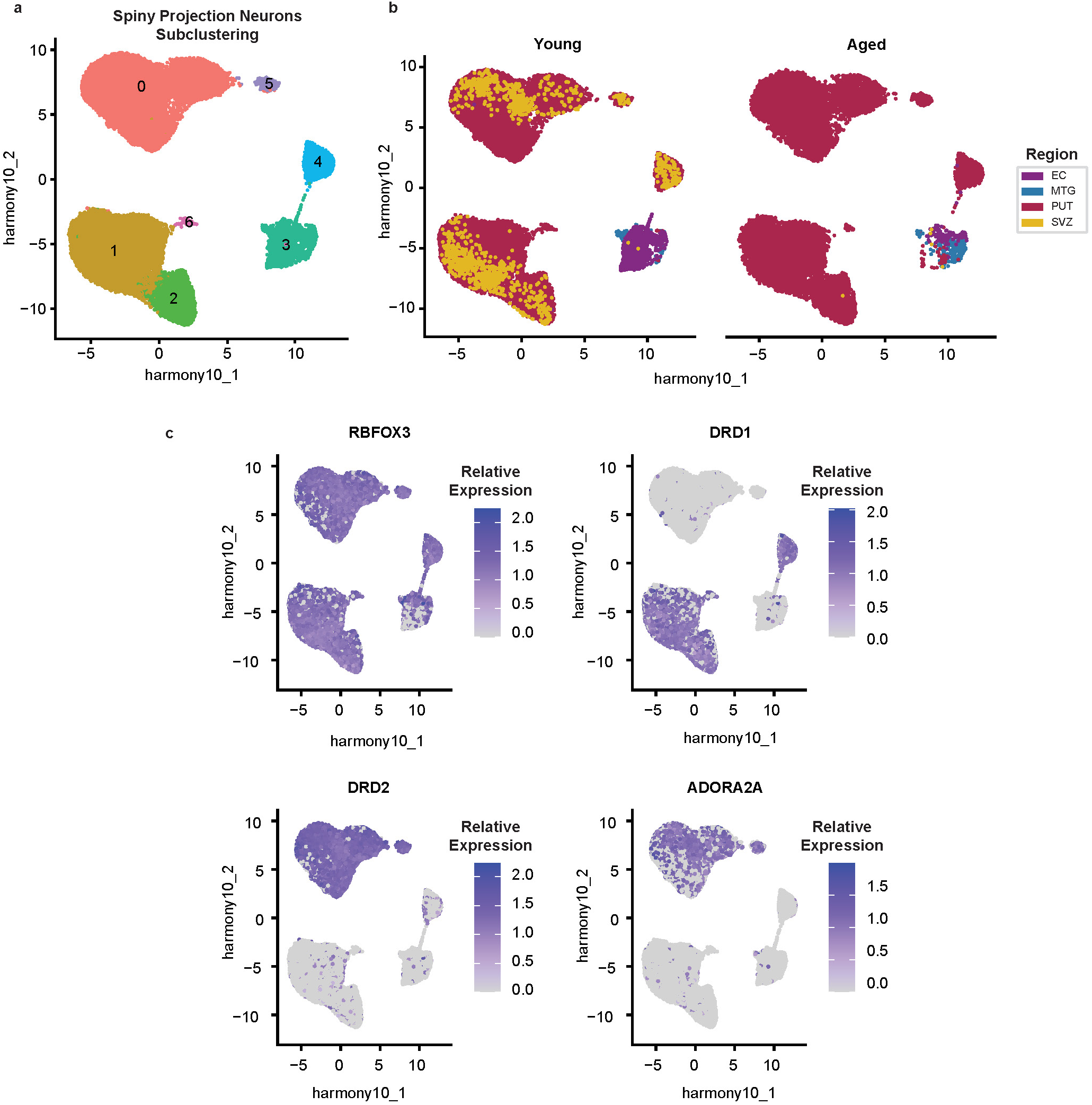
Subclustering of nuclei annotated as spiny projection neurons affirms regional specificity for putamen and indicates probable misannotated nuclei. **(a)** Subclustering of nuclei annotated as spiny projection neurons in the initial round of clustering results in 7 sub-clusters. **(b)** Separating these nuclei by age group (young, left; aged, right) and coloring them by brain region of origin reveals that subclusters (0-2 and 5-6) largely come from the putamen. Nuclei from other regions were few in number (n=3,545), so they were re-annotated as “Other” and excluded from further analysis. (**c**) Relative normalized expression levels for known SPN marker genes differentiate subclusters and confirm SPN identity.

**Supplemental Figure 4:**
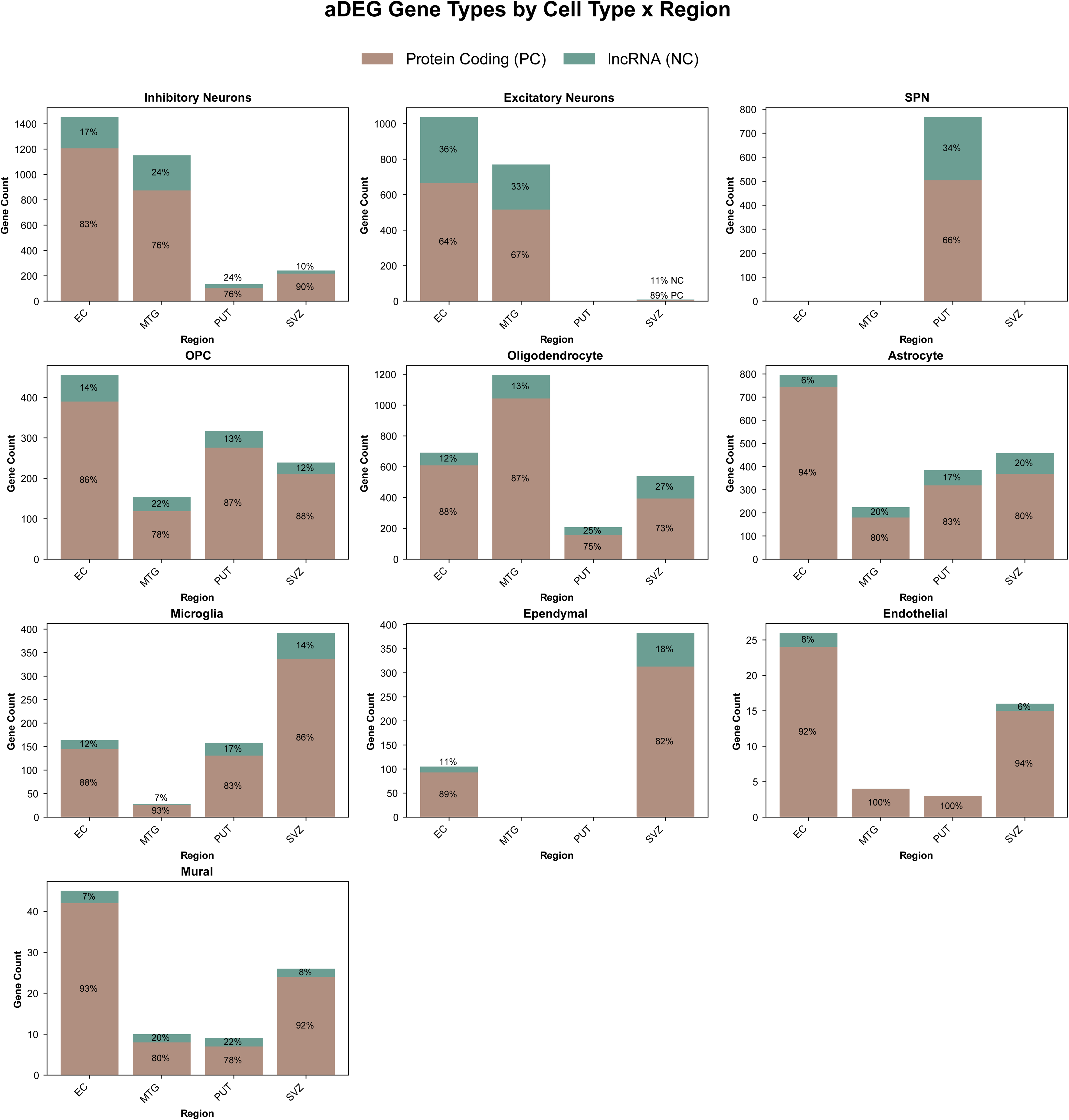
Majority of aDEGs are protein coding across cell-types and regions. Classification of aDEGs as protein coding or long non-coding RNAs (annotations associated with refdata-gex-GRCh38-2020-A) across all cell-type-region combinations reveals that the majority of aDEGs are protein coding.

**Supplemental Figure 5:**
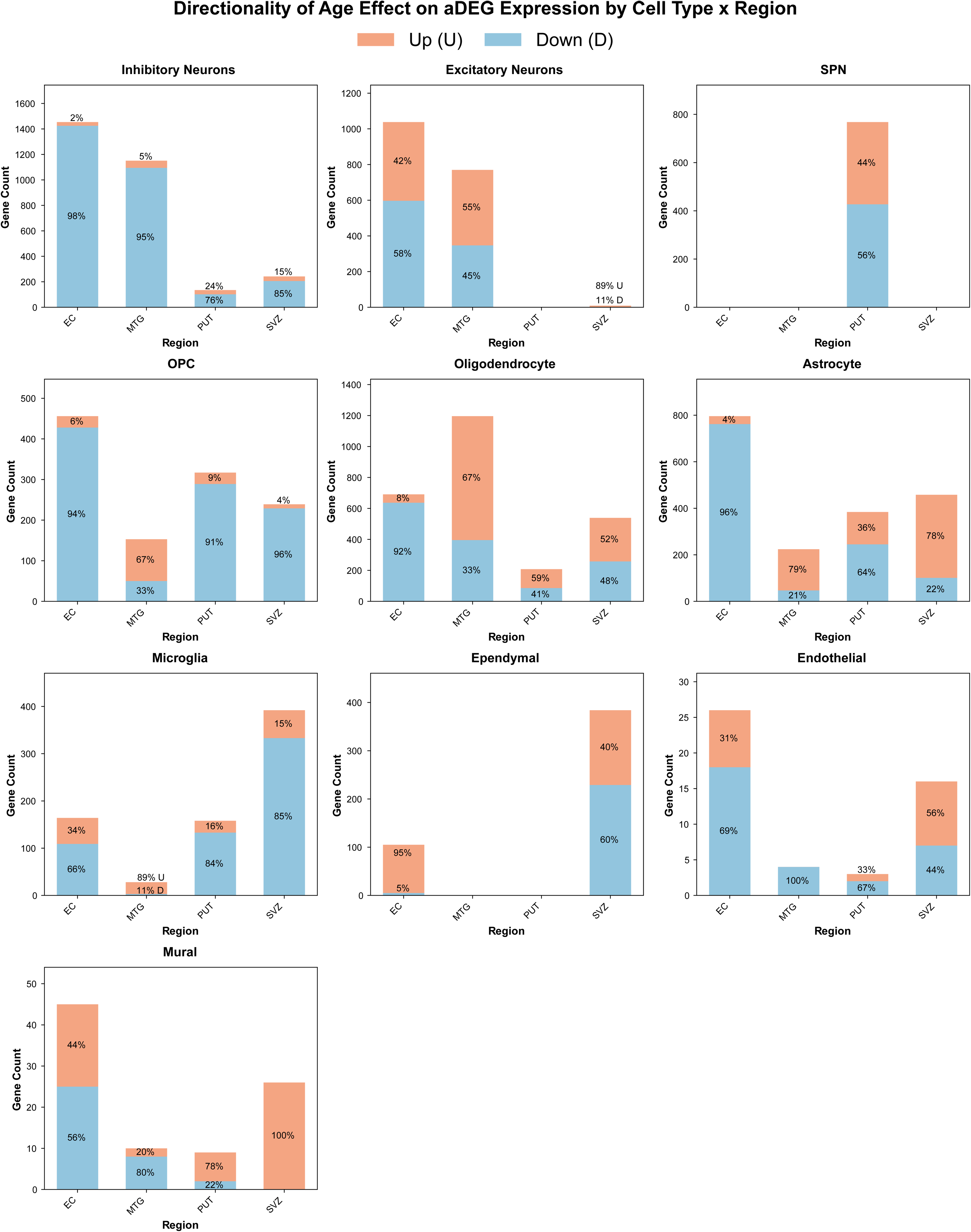
aDEG expression levels tend to decrease with age across most cell-type by region combinations. aDEG expression direction (i.e. increase or decrease) across all cell-type-region combinations suggests that overall, the majority of aDEGs tend to decrease in expression with age.

**Supplemental Figure 6:**
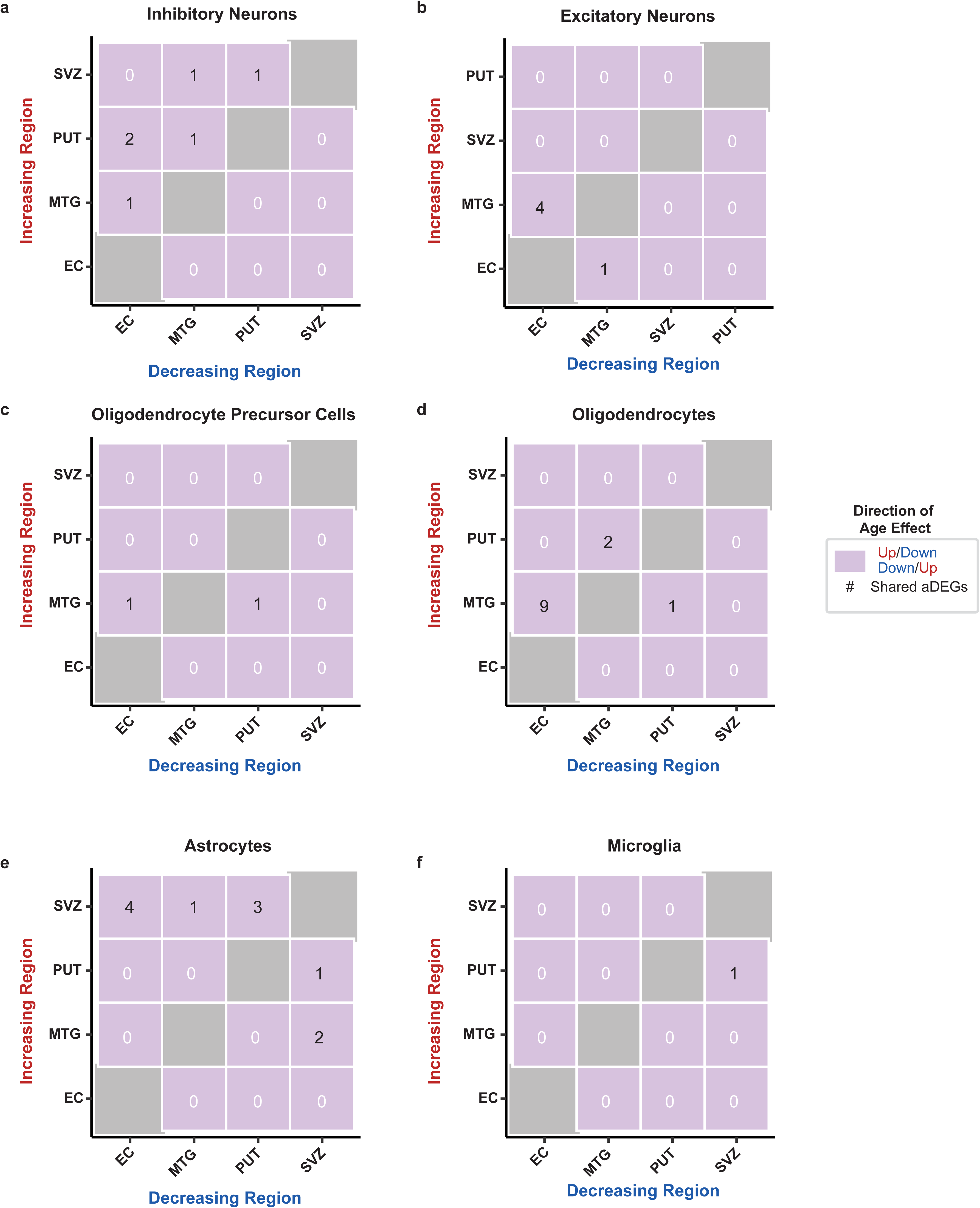
Few regionally-shared aDEGs with opposing age-effect directions across cell-types. Pairwise comparison of shared aDEGs that differ in age-effect direction between regions (discordant) within a given cell-type reveals few instances of discordant aDEGs.

**Supplemental Figure 7:**
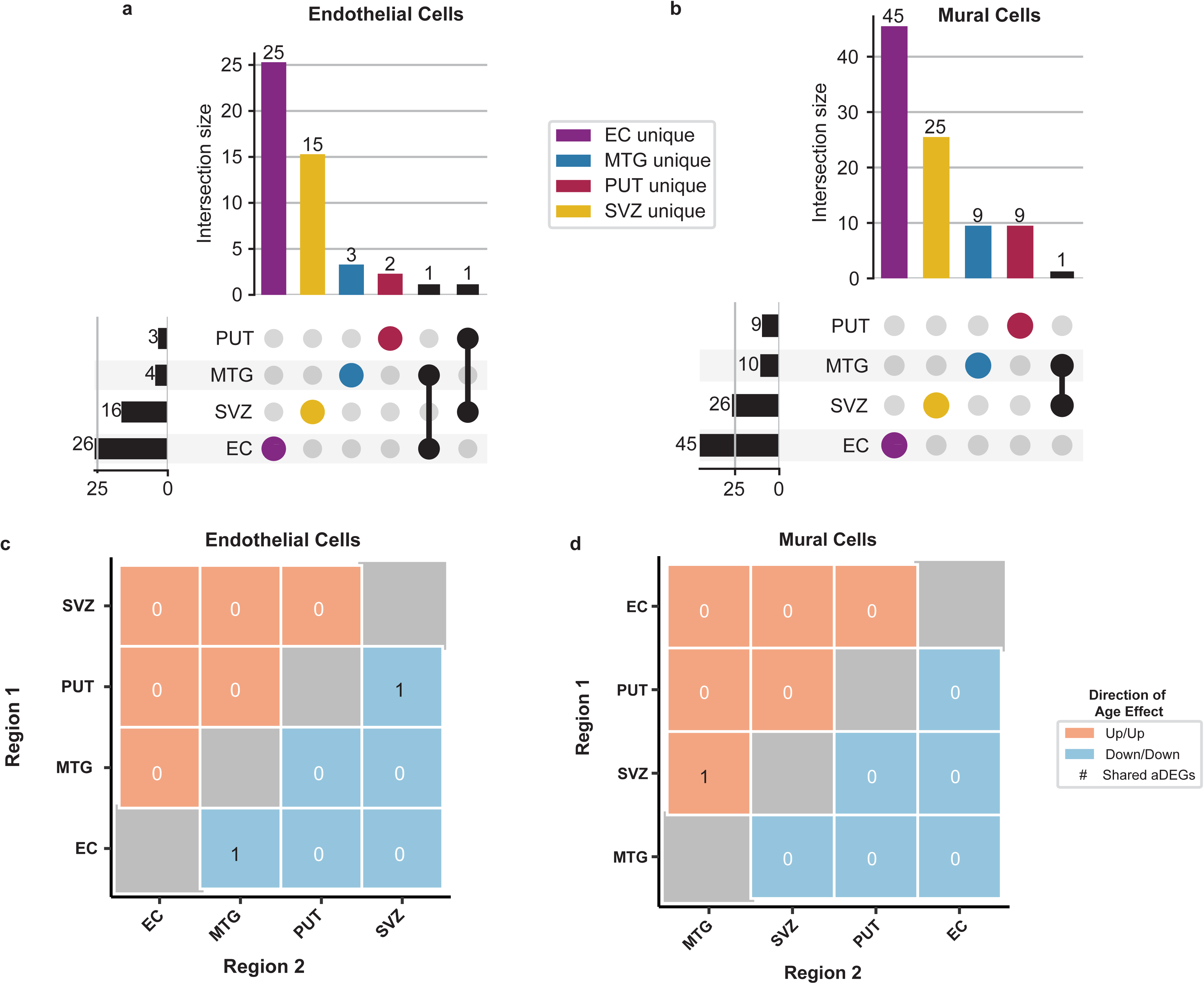
Pericytes exhibit few aDEGs with the majority being in the EC and SVZ. Regional distribution and overlap of pericyte cell-type aDEGs indicate that the majority of aDEGs are unique to a particular region. Both **(a)** endothelial cells and **(b)** mural cells have the majority of aDEGs localized to the EC followed by the SVZ. Pairwise comparison of shared aDEGs within pericytes suggests little regional sharing within both cell-types. Heatmap values representing count of aDEGs indicate the number of aDEGs changing in the same direction (concordance)--either increasing (positive, red) or decreasing (negative, blue)–in both of the indicated regions. **(c)** Endothelial cells showed 2 negatively concordant aDEGs one between the EC and MTG and the other between the PUT and SVZ. **(d)** Mural cells showed 1 positively concordant aDEG between the SVZ and MTG.

